# Red deer individual landscapes of fear in response to human recreation

**DOI:** 10.1101/2024.06.20.599860

**Authors:** Bjoern Boehme, Anne Peters, Veronika Mitterwallner, Fabian Sommer, Marco Heurich, Anne Gabriela Hertel

## Abstract

Animals adjust behavior to changes in perceived predation risk, even when risk is non-consumptive, as is the case for human recreation. However, individuals within populations can differ greatly in their plasticity towards perceived risk, especially when antipredator responses incur fitness costs via lost foraging opportunities. Therefore, risk-benefit trade-offs are usually made at the individual level. We test whether red deer (*Cervus elaphus*) inhabiting the Bavarian Forest National Park show individual variation in behavioral plasticity towards human disturbance. We measured human disturbance as the number of recreationists on the closest trail on a given day and used random regression to dissolve movement responses, measured as hourly step-length, of 63 GPS-collared red deer females to fine scale spatio-temporal variation in disturbance. We quantified behavioral state specific between-individual variation along the disturbance gradient. At the population level, red deer responded to recreational disturbance during the middle of the day only, and reduced movement in response to recreation. However, population patterns masked strong between-individual variation in plasticity to recreation activity during the morning, midday, and evening, such that some deer reacted with flight while others responded little. Behavioral responses were mediated by vegetation cover with stronger responses in more open areas. Flight responses are costly due to lost foraging opportunities. A shift in risk-benefit trade-offs when risk is non-lethal, as is the case for recreation in many national parks, may favor more human-tolerant individuals over time.

**Lay summary:** We assessed the individual response of red deer towards recreational activities in the Bavarian Forest National Park, using GPS-telemetry data and modelled daily numbers of tourists on trails. Red deer varied in their fear of humans, most red deer adjusted movement on days with many park visitors, but some individuals did not respond to human disturbance. Individual variation in movement increased with human disturbance. Foraging-risk trade-offs may promote behavioral variation of red deer and their tolerance of humans.

## 1. Introduction

Animals make state-dependent decisions about when and where to move and dynamically adjust movement to the local environmental conditions (Nathan et al., 2008). For example, movement decisions differ when animals are foraging versus selecting resting sites and movements often slow down at steeper slopes and denser vegetation. When animals are active, their movement decisions are dominated by two main drivers: finding areas of high forage availability and avoiding predation risk (Nathan *et al*. 2008).

How animals manage this trade-off has been at the heart of ecological research, referred to as the “landscape of fear”. Landscapes of fear describe where and when prey perceive predation risk as high and adjust their behavior accordingly (Gaynor et al., 2019; Kohl et al., 2018). Likewise to natural predators, human activities contribute to the landscape of fear and causes behavioral responses in wildlife that resemble anti-predator behavior (Frid and Dill, 2002). While wildlife have developed responses towards potentially threatening stimuli in general to increase survival, humans act as superpredators on all wildlife species and their consumptive and non-consumptive effects far exceed those of natural predators in many systems (Clinchy et al., 2016; Darimont et al., 2015; Zanette et al., 2023). Changes in wildlife behavior, i.e., behavioral plasticity, is expected to increase exponentially with perceived risk (Lima and Dill, 1990). However, when the variation in risk is spatially or temporally predictable, the risk allocation hypothesis predicts that wildlife shows disproportionately stronger antipredator responses during times or in places of highest predation risk in order to maximizing foraging opportunities (Creel et al., 2005; Ferrari et al., 2009; Gaynor et al., 2019). For ungulates, foraging and risk avoidance (sometimes of multiple predator species, Lone et al., 2014) leads to trade-offs as the highest quality food can often be found in open meadows where visibility and predation risk are higher (Mao et al., 2005).

### Individual landscapes of fear

Recent advances indicate that prey responsiveness to perceived landscapes of fear, expressed for example as plastic adjustments of movements to increasing risk or infrastructure, may vary substantially among individuals, with some individuals adjusting behavior more strongly to perceived risk than others (Dammhahn and Almeling, 2012; Dammhahn et al., 2022; but see Salazar et al., 2023). This is also true for landscapes of fear created by humans (Found and St. Clair, 2018; Xu et al., 2023). This creates an interesting situation where some individuals may accept even high perceived mortality risks more readily at the potential benefit of increased forage intake and higher reproductive fitness (Dammhahn et al., 2018). Contrarily, comparative studies under controlled conditions suggest that high perceived predation risk may limit the expression of between-individual variation and force individuals to conform to a common behavioral phenotype, while behavioral variation can be expressed more freely under low perceived risk (Dingemanse et al., 2009). However, under controlled conditions, risk-benefit trade-offs are solely hypothetical as selection pressures are absent (i.e. no fitness benefits of higher resource acquisition and no mortality cost of risky behavior) and therefore laboratory findings may not apply in the wild (Moiron et al., 2020). Whether individuals differ in behavioral plasticity to perceived risk and, in particular, whether individual variation is promoted or restricted at higher levels of risk is therefore a question in behavioral ecology and wildlife biology that warrants further investigation.

### Recreation as source of human disturbance

While natural predators always incur a real predation risk, and therefore warrant antipredator behavior (at least when spatio-temporal risk is high, Ferrari et al., 2009), humans are a special case. The fear of the human predator is ubiquitous in wildlife, including apex predators (Zanette et al. 2023), and has likely been engrained through centuries of heavy harvesting and strong selection of fearful individuals (Penteriani et al., 2016; Zedrosser et al., 2011). Therefore, wildlife also adapts behavior to non-lethal recreational activities (Taylor and Knight, 2003). Many studies have demonstrated a behavioral response of wildlife to recreational activities, including both increases or decreases in movement (Ladle et al., 2019; Neumann et al., 2010; Olson et al., 2018; Ordiz et al., 2013). The specific behavioral response (freeze or flight) may particularly depend on the spatial context, with wildlife choosing the strategy that will most likely decrease the risk of encountering humans. If wildlife can move to disturbance free areas, increasing their movement rate may be a suitable response (Neumann et al., 2010). However, if disturbance free areas are not readily available, decreasing movement and selecting dense vegetation may be the more suitable strategy as dense vegetation can provide refuge from human activities and decrease the perceived risk (Jayakody et al., 2008; Olson et al., 2018; Ordiz et al., 2013; Versluijs et al., 2022). High human disturbances by recreational activity, however, may lead to wildlife generally avoiding to be active during times when humans are active (Marchand et al., 2014). Human recreation is particularly high in many national parks, areas that are designated to protect nature and provide refuge for wildlife. At the same time, an integral goal of national parks is to promote recreation and environmental education within their area. In fact, nature tourism and wildlife viewing are becoming important agents to increase the public’s appreciation for protecting nature (Böhn, 2021). Understanding the impact of recreational activities on wildlife and implementing measures to improve the visitors’ chances to experience wild spaces, may therefore be a useful tool to support conservation efforts and public acceptance of protected areas (Marion et al., 2016).

We here aim to test whether ungulates differ in behavioral plasticity towards gradients of perceived risk incurred by recreational activities in a national park. We are particularly interested in whether fear of humans promotes or restricts between-individual variation when the perceived risk is high compared to contexts that are perceived as less risky. We use a model system of an ungulate prey, the red deer (*Cervus elaphus*), responding to spatio-temporal variation in human presence, their main predator: Red deer have been hunted to extinction throughout large parts of Central Europe, but have since recovered in isolated populations and are nowadays again one of the top game species. Our study area, the Bavarian Forest National Park (BFNP) offers a unique situation of a large and connected non-hunting zone, which was first implemented 25 years ago in 1997, representing approximately five red deer generations (Coulson et al., 1998). The non-hunting zone nowadays encompasses 75% (18,700 ha) of the park area and further extends into the non-hunting zone of the neighboring Šumava National Park. However, recreational activities can be carried out in most of the BFNP area, even if restricted to the national park’s trail system, with the most intense frequentation by recreationists over the summer months (Porst et al., 2022). Recreational activities on trails have been constantly monitored in the BFNP using wildlife cameras, originally deployed in the frame of a large carnivore monitoring along trails. These cameras formed the basis of a model predicting daily number of recreationists on every official trail in the park (Mitterwallner et al.). However, as suggested previously, spatial context is expected to modulate the impact of human disturbance on red deer behavior. For example, deer may be less responsive to humans when hidden in dense vegetation. In the BFNP, the availability of fine scale Lidar scans provide the basis for a high-resolution shrub and visibility layer, allowing to control for the spatial context of concealing vegetation cover (Zong et al., 2021). Combining the detailed information on recreational activities and available cover with red deer GPS-movement data collected from 2018 until 2021, our model system provides a natural experiment to study spatio-temporal dynamics of prey responses to perceived risk and to test the synergetic effects of human disturbance and cover on the expression of between-individual variation in red deer movement behavior.

Our main objective for this study was to determine whether human disturbance promotes or restricts individual variation in red deer movements. Theory predicts that individual variation could either be promoted (Dammhahn et al. 2022) or buffered (Dingemanse et al 2009) at higher levels of disturbance, depending on whether risk-benefit trade-offs are incurred. We hypothesized, H1) that female red deer adjust movement in response to recreational activity in their vicinity. Specifically, we predict that red deer increase movement as a flight response to move to less disturbed areas, especially when concealing vegetation cover is low. However, flight responses bear a foraging cost and human recreation is temporally highest during the daylight hours. In line with the risk allocation hypothesis we therefore predict that red deer respond strongest to the number of daily recreationists during the daylight hours. In addition, more risky individuals (i.e., those that respond less to human disturbance) may have a fitness benefit because they can access more or better forage compared to responsive individuals. We therefore hypothesize H2) that female red deer in the BFNP will display individual variation in behavioral plasticity to human recreational activities. We predict that between-individual variation increases with perceived risk, such that some individuals respond with increasing movement to higher perceived risk, while others do not adjust movement. We further predict that available cover will reduce the perceived disturbance, hence red deer in areas with higher cover will show less between individual variation in movement responses as recreational disturbances increase, than in areas of low cover.

## 2 Methods

### 2.1 Study area and species

The study took place in the Bavarian Forest National Park (BFNP, 49°3’19’’N; 13°12’9’’E, 254 km^2^), Germany’s oldest national park. The forest has been largely unmanaged for the past 50 years creating, together with the Czech Sumava NP, the largest strictly protected forested area in Central Europe (Heurich et al., 2015). More than 60% of the area is covered by coniferous forest (mainly Norway spruce, *Picea abies*), 20% by mixed forests, and 14% by grasslands. The area is characterized by low to intermediate elevations (500 - 1453 m.a.sl). Nowadays the BFNP is a popular area for recreational activities with about 300 km of hiking and 150 km of biking trails and approximately 1.3 million visitors annually. Several access points with designated parking areas provide access to a dense network of hiking trails. The park is home to a thriving red deer population, which was formerly extinct in the area, alongside roe deer (*Capreolus capreolus*) and wild boar (*Sus scrofa*). Natural predators are the Eurasian lynx (*Lynx lynx*), feeding however mostly on calves (Belotti et al., 2015). Grey wolves (*Canis lupus*) have been recolonizing the area since 2015 (Hulva et al., 2024), but wolf presence was very low in most of the BFNP area until 2023, leading to low or missing predation risk by wolves during previous years. The red deer population is partially migratory with about 55% of deer remaining in their summer ranges (Peters et al., 2019). Hunting is strictly prohibited throughout 75% of the study area in the BFNP. However, the red deer population is still closely managed. Annual quotas are determined based on counts, quotas in previous years and browsing damage. Hunting is allowed from June to January in the park’s management zone (25% of the study area in the BFNP) and, which is located along its borders and outside the park’s boundaries where red deer are mainly hunted from high stands. Additionally, red deer are culled in winter enclosures. These structures are large fenced areas including feeding stations, intending to reduce damages in managed forests near the border of the park as red deer would migrate to lower elevations and thereby out of the BFNP when food becomes scarce due to snow cover in winter (Möst et al., 2015).

### 2.2 Red deer movement data

Between 2018-2021 we captured and collared 63 mature female red deer in four winter enclosures. While most individuals were equipped with collars without chemical immobilization, 13 individuals were immobilized via teleinjection, with the Hellabrunner mixture (Ketamin and Xylazine). Experimental procedures were approved by the Ethics Committee of the Government of Upper Bavaria (permit number: ROB-55.2-2532.Vet_02-19-130; ROB-55.2-2532.Vet_02-17-190). Red deer were fitted with Vertex Plus GPS collars from Vectronic Aerospace GmbH, scheduled to record a GPS location every hour. We removed positions resulting in unusually long displacements (> 4000 m) or unusually high speeds (> 4.17 m /s). We included locations from the beginning of May to the end of October, i.e., the period of the year when red deer are not in the winter enclosures. Most individuals were followed for more than one year (range 1 – 4 years) resulting in 104 monitoring years (deer-year). We constructed individual movement trajectories and calculated step-lengths using the R package amt (Signer et al., 2019). We used hourly step length, i.e., the distance moved between two consecutive locations, as a measure of responsiveness to perceived risk because it can be readily interpreted in the framework of a flight response, it is robust to outliers and it measures behavior at the finest temporal scale possible given the programmed GPS interval (1 hour). Step-length was calculated as the Euclidean distance between successive, hourly positions of an individual. We extracted spatial covariates for the leading GPS location of each step-length. After cleaning, a total of 273,124 successful GPS locations were taken forward into analysis. Of these positions, 85% (231,455 positions) were located within the park’s core area where hunting is prohibited while 15% (41,669 positions) fell into the management zone where hunting is allowed.

### 2.3 Spatial covariates

#### 2.3.1 Recreational disturbance

From November 2020 – November 2021, 61 wildlife cameras were distributed along the BFNP trail network in the frame of the large carnivore monitoring, providing additional information on recreation activities on trails at any given day of the year. In a total of 24,461 camera days, we captured > 700,000 pictures. We used the modelling approach from (Mitterwallner et al. under review) to model the number of recreationists on a daily basis on a given hiking trail and to predict spatio-temporally explicit trail use by recreationists. Models included information about weather, weekend and holiday seasons and the location of the path relative to parking sites and huts, the density of parking sites and touristic points of interest in the area as well as information on the path access restrictions. The models explained 44/47% of the variance in the number of daily recreationists We extracted the closest path segment for each red deer GPS-location and predicted the number of visitors expected on this path on the given day on which the red deer GPS-location was recorded.

#### 2.3.2 Visibility at 140cm from LiDAR data

In June 2017, the entire national park was subject to airborne LiDAR scanning (ALS) from a helicopter using a Riegl LMSQ 680i instrument. Scans were conducted with a helicopter from 550m flight altitude. The average point density was 30 points/m2 and the vertical and horizontal accuracy of the ALS data was 6 and 5 cm (Krzystek et al., 2020). Fine scale terrestrial LiDAR scans at 93 forest sampling plots were used to build a random forest model that best predicted visibility (Zong et al., 2021, 2022). Using ALS sensed metrics of the five covariates best predicting visibility, visibility was predicted across the whole park. These five covariates were: 10th height percentiles of canopy returns, 70th height percentiles of understory returns, normalized relative point density (NRD), percentage of NRD and coefficient of variation of 99th height percentiles of understory returns. Habitat visibility was specifically predicted at 140 cm height (equivalent to a standing height of red deer). The resulting visibility layer had a spatial resolution of 35m. Please refer to (Zong et al., 2021, 2022) for methodological details.

#### 2.3.3 Elevation and temperature

We extracted elevation above sea level from a raster layer at a 25 m resolution. We further used the chillR R package (Luedeling et al., 2023) to extract daily temperature data (mean and maximum daily temperature) from the closest weather station, situated in Churanov, about 18km from the centroid of all red deer GPS data on the Czech side of the Bavarian Forest – Sumava national park.

### 2.4 Statistical analyses

The main aim of this study was to assess between-individual variation in plasticity towards human disturbance with random regression. Movement trajectories are a great source of data providing many repeated measures of an individual’s behavior (here step length) along ecological gradients. However, movement data are inherently spatially and temporally autocorrelated, particularly when fix-intervals are short, under limited movement activity (i.e., stationary behavior), or when the spatial resolution at which ecological covariates are measured is coarser than the step length. To avoid pseudoreplication and, thereby upward bias of between-individual variation estimates. (successive records of an individual reflecting the same behavior under the same conditions, Niemelä and Dingemanse, 2017), we fit random regression models (2.4.2) to specific times of day, corresponding to distinct behavioral states of peak activity and resting (2.4.1). To this end we first determined month specific red deer diel activity (2.4.1) patterns to then select step-length measures at discrete times of day, corresponding to distinct behavioral states, for further analyses. The response variable in the random regression models (2.4.2), our main analysis, was therefore one daily measure of hourly step length per individual at the specific hour of the day. With this approach we effectively ruled out any temporal autocorrelation causing pseudoreplication (successive records of an individual reflecting the same behavior under the same conditions) while any remaining temporal autocorrelation (individuals behaving in a similar manner on successive days) is biologically informative to our research question, i.e., whether individuals behave consistently over time.

#### 2.4.1 Diel activity patterns

We determined diel patterns of movement activity, measured as step length, by fitting a cubic spline over the time of day. We accounted for between-individual variation in activity patterns with a random intercept for each deer-year and random slopes over time of day. Diel activity and between-individual variation therein may shift over the summer due to changing daylight or disturbance, however, further nesting of cubic spline random slopes was not possible. We therefore fitted individual models for each month (May to October, six models). We log-transformed step lengths as they were highly skewed towards shorter distances. We quantified and visually examined whether individual variation in diel activity patterns existed. We then inspected the population average diel activity profile and selected the month-specific hours of the day when activity peaked or was at its lowest for further analysis.

#### 2.4.2 Random regression

Given the crepuscular bimodal activity pattern, we fitted four behavioral state specific models (Table 1), one for the respective (month specific) hours of the day when activity peaked or troughed. We modelled step length (log-transformed) as a function of human disturbance interacting with visibility at 140cm height. Red deer were mostly in close vicinity to the next trail where the disturbance was measured (median distance 155m, few positions up to 2km from the closest trail), owing to the high trail density in the park. Elsewhere, the effect of recreational activity of mule deer behavior dropped beyond 390m from a trail (Taylor and Knight, 2003), however, our study area consists mainly of forest, providing more vegetational cover than the study area by Taylor and Knight. We therefore only included positions no farther than 300m from a trail (80% of all red deer positions in the original dataset), which we presumed to be within the perceptional range of red deer. The four models were therefore fit on 10,490 - 12,246 individual, day, and time specific hourly step lengths. We expected that visibility, i.e., cover, would moderate the effect of human disturbance on red deer behavior and therefore fitted an interaction between the two terms. We further accounted for known effects of elevation (nonlinear second-order polynomial), canopy cover, and maximum daily temperature on movement behavior. Last, we fitted random intercepts for each deer-year and random slopes over human disturbance interacting with visibility to account for between-individual variation in behavior plasticity. To interpret effects, we predicted red deer movements from low to high human disturbance (resp., 1 – 40 recreationists on the closest trail, Supplementary material Figure S1) and at low and high visibility (15^th^ and 85^th^ percentile of visibility distribution, Figure S2), while we assumed an average elevation and canopy cover.

**Table 1.**
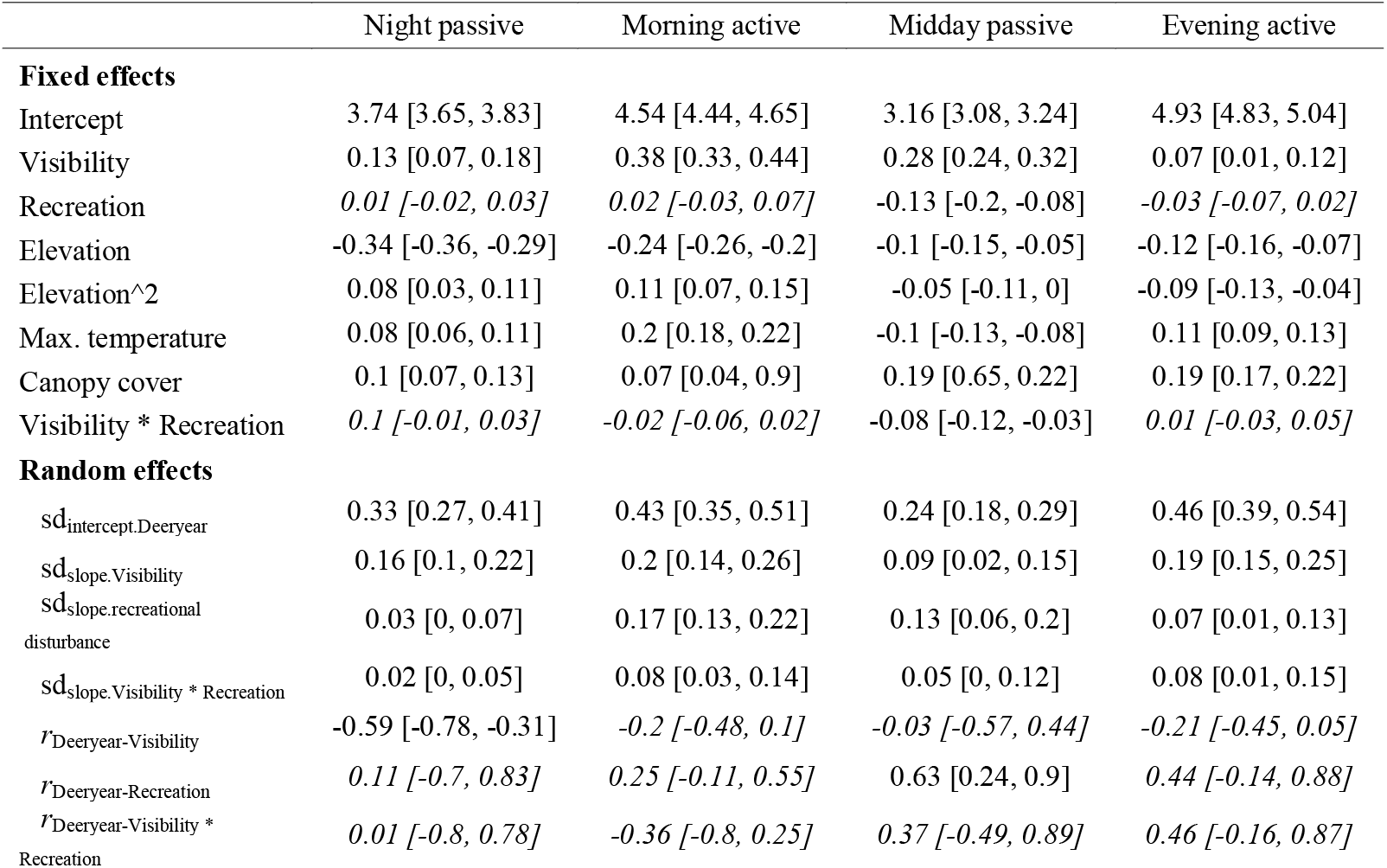
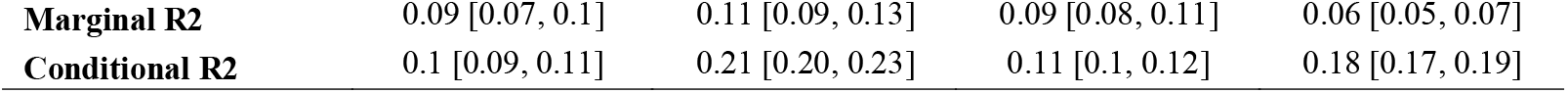
Estimates and 95% credible intervals (in parentheses) of random regression models at four behavioral state specific times of day: night passive, peak morning activity, midday passive and peak evening activity. The variance explained the fixed effects (marginal R2) and by the combined fixed and random effects (conditional R2) is given under each model.

#### 2.4.3 Between-individual variation in plasticity

We assessed between-individual variation in plasticity using estimates of conditional repeatabilities along the gradients of human disturbance and visibility following (Schielzeth and Nakagawa, 2022). Conditional repeatability is the ratio of between-individual variance to phenotypic variance (between-individual variance plus residual variance) at specific levels along the environmental, i.e., random slopes gradient. The index ranges from 0 to 1 with higher values indicating greater between-individual behavioral variability. Across a broad suite of behaviors, the meta analytical mean repeatability of behavior has previously been estimated at 0.37, while repeatability values < 0.2 can be considered low (Bell et al., 2009). For interpretation, we consider repeatability values > 0.2 as moderate and repeatability values > 0.37 as high.

All random regression and diel activity models were fit with a Gaussian family on log transformed step length using the R package brms (Bürkner, 2017). We used uninformed default flat priors for fixed effects and student-t priors for random effects. We ran four chains over 2000 iterations with a warmup of 1000 and a thinning interval of 2. The model inference was based on 2000 posterior samples and had satisfactory convergence diagnostics with ^R < 1.01 and effective sample sizes > 1000. Posterior predictive checks recreated the underlying Gaussian distribution well (Figure S3).

## 3. Results

### 3.1 Pronounced crepuscular activity pattern with little between-individual variation

Red deer showed a crepuscular activity pattern over the entire summer, which shifted with changes in sunrise and sunset (**Fig 1**). Movement distances were longest during the morning and evening twilight. Movement distances were shorter during the night and daylight hours with pronounced resting during midday. Although between-individual variation in movement distances were evident, we found little between-individual variation in the shape of the diel activity patterns, i.e., all individuals followed a bimodal activity pattern. We therefore proceeded to analyze between-individual variation in movement during the month-specific hours of highest and lowest activity using random regression (gray bars in **Fig 1**).

**Fig. 1.**
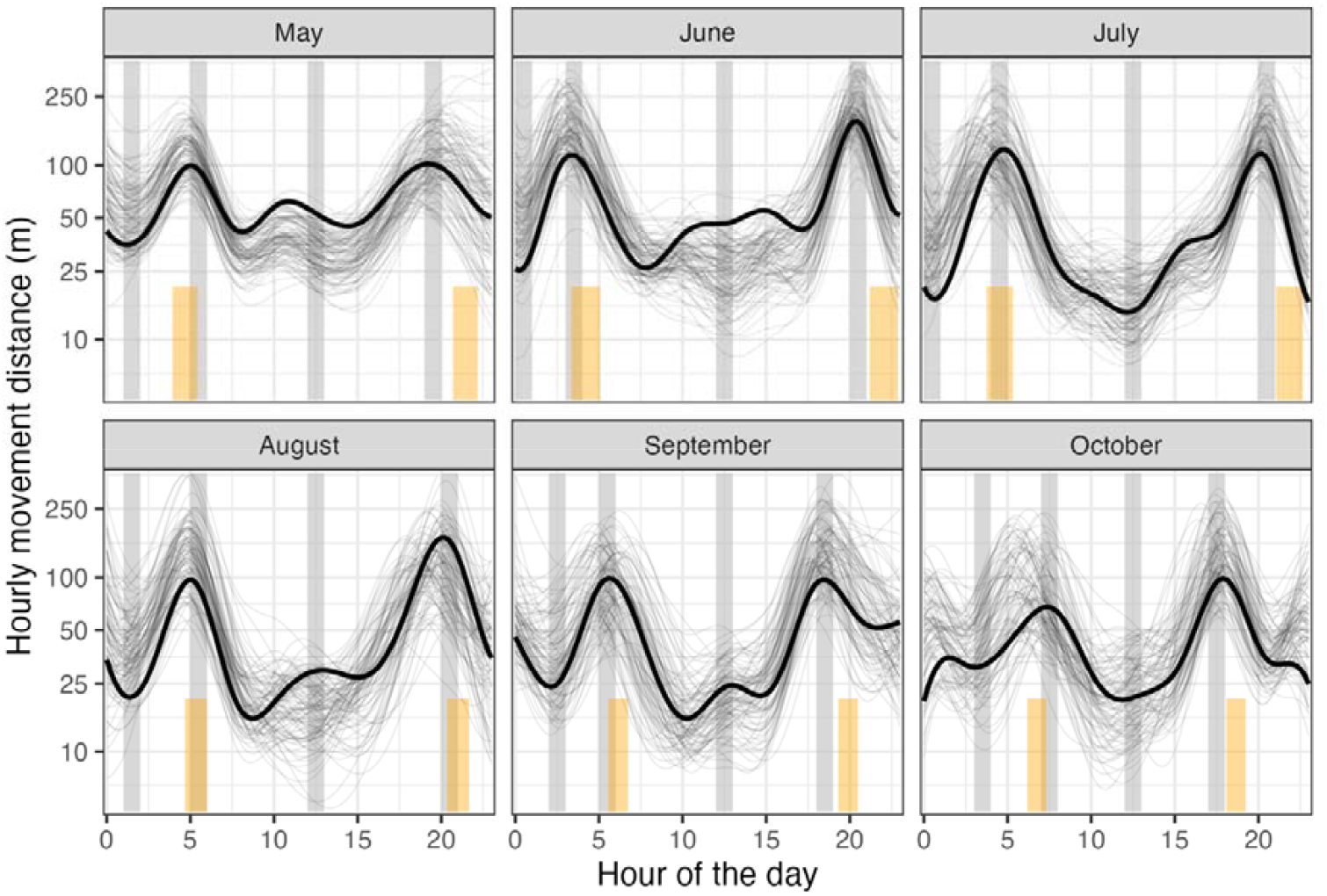
Population (thick line) and individual level (thin lines) diel activity patterns of red deer monitored in the Bavarian Forest National Park from May to October. Orange bars indicate times of dusk and dawn (from nautical dawn until sunrise and from sunset to nautical dusk). Grey bars indicate hours of peak activity and lowest activity. We took forward daily movement distances recorded at hours of highest and lowest activity to model between-individual variation in red deer responsiveness towards human disturbance.

### 3.2 Population level responses to recreational disturbance and visibility

Selecting the month-specific hours of highest and lowest activity we fitted four behavioral state specific models using step length measures from 104 deer monitoring years: a night passive model (based on 12,246 step length measures at 12 – 3 am, depending on the month, see **Fig. 1**), a morning active model (based on 12,237 step length measures between 3 – 7 am, **Fig. 1**), a midday passive model (based on 10,490 step length measures at 12 pm, **Fig. 1**), and an evening active model (based on 11,406 step length measures between 6 – 8 pm, **Fig. 1**). Recreational disturbance on trails varied from 1 to 111 recreationists per day but red deer mostly experienced low numbers of recreationists per day (1^st^ – 3^rd^ quartile = 5 – 11, **Fig. S1**), though almost all red deer were exposed to a moderate number of 20 recreationists at some point (**Fig. S1**). Red deer were primarily found in intermediately dense vegetation with moderately high visibility, except for the middle of the day when they were found in areas with lower visibility (**Fig. S2**). Maximum daily temperature ranged from 0°C to 29°C (1^st^ – 3^rd^ quartile = 11°C – 20°C) and elevation from 628 – 1374 m.

Behavioral responses of red deer to recreational disturbance were context and behavioral state dependent, both at the population and the between-individual level (**Table 1, Fig 2**). On the population level, red deer responded to increasing numbers of recreationists during the middle of the day (midday passive) but not during the night passive, morning active, or evening active bout (partial support for recreational disturbance term, **Table 1**). Red deer moved over longer step lengths in open than in dense vegetation at all times of day (support for visibility term, **Table 1**) but particularly during the morning active and midday passive bout. Additionally, vegetation density moderated the effect of recreational disturbance on red deer movement during the middle of the day (support for the interaction term, **Table 1**). While, in line with their diel activity patterns, red deer generally moved very little during the middle of the day resting bout, they further reduced movement from an average of 44 m/hr to 12 m/hr on days when disturbance was high and when in open vegetation (**Fig 2**). Red deer movements were additionally governed by elevation and canopy cover, with shorter step lengths at higher elevation, and longer step-lengths in areas with denser canopy cover (**Table 1**). With increasing maximum daily temperature, red deer moved over shorter step lengths in the middle of the day while increasing movements at all other times of day.

**Fig 2.**
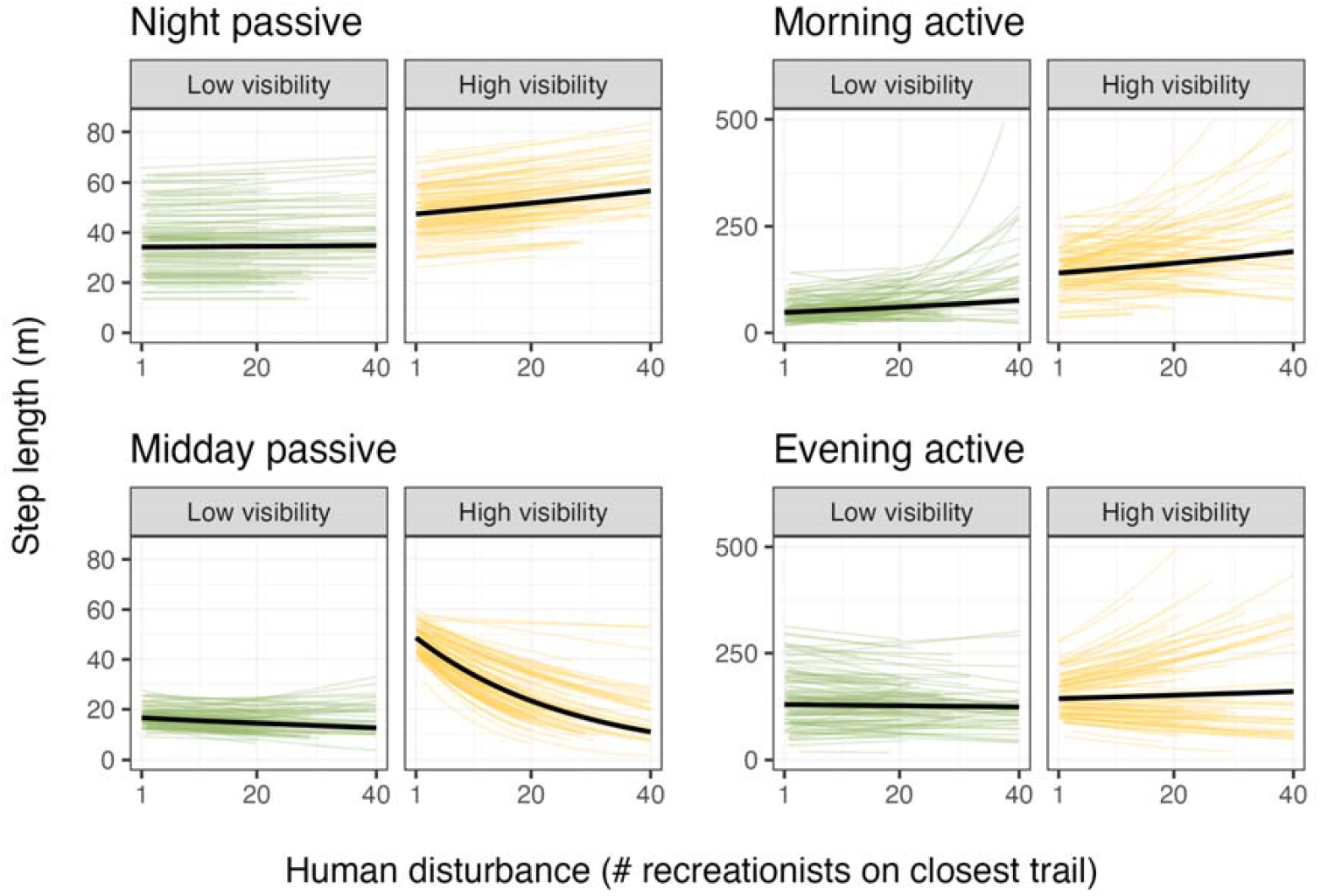
Model predictions of the effects of recreational disturbance and vegetation cover on movement of red deer in the Bavarian Forest National Park during peaks and troughs of daily activity. Population level effects are shown as a black line with individual-level random intercepts and slopes in green (low visibility) and yellow (high visibility). Low and high visibility were fixed at the 15^th^ and 85^th^ percentile of the observed distribution of cover at red deer locations. Individual predictions extend only over the disturbance gradient experienced by each individual. recreational disturbance affected the movement of red deer most strongly during the morning active and midday passive bout. Effects are back transformed to the linear scale for ease of interpretation, mind the different y-axes extents for movement during passive (left) and active (right) bouts. Y axes are truncated to a maximum of 500m for better readability, although several individuals increased movement up to 1900m at very high recreational disturbance.

### 3.3 Between-individual variation in responsiveness to recreational disturbance

Red deer varied most strongly in their responsiveness towards recreational disturbance during the morning bout of activity (**Table 1**). For example, under high visibility the most responsive individual increased movement fourfold, from 296m/hr under low disturbance to 742m/hr under high disturbance (**Fig 2**), while the least responsive individual only increased movement by 20m/hr (from 205m/hr to 224m/hr). Individual variation in red deer movement was therefore mainly expressed when disturbance was high with individuals reacting differentially responsive to increases in perceived risk. During the morning activity bout, there was no correlation between an individual’s expression at low and high recreational disturbance, i.e., deer that moved more or less under low disturbance were equally likely to respond more strongly to recreational disturbance. Consequently, mean conditional repeatability in the morning (**Fig 3**), indicative of the amount of between individual variation in movement, increased from 0.16 under low disturbance to 0.5 under high disturbance. As shown in section 3.1 and 3.2, on the population level, red deer move little in the middle of the day but further decreased movement on days of high disturbance, however some individuals did not reduce movement in response to recreational disturbance (**Fig 2**): the most responsive individual decreased movement from 40 to 2 meter, while the least responsive individual maintained an average hourly step length of 50 meters. In fact, the positive correlation between random intercept and slope (**Table 1**) shows that those individuals which already move more during the middle of the day when recreational disturbance is low, were more likely to maintain movement with increasing disturbance, while red deer that move over shorter distances to begin with decreased movement more strongly. Individual variation in red deer movement in the middle of the day therefore also increased with recreational disturbance (from conditional repeatability = 0.03 at low disturbance to conditional repeatability = 0.3 at high disturbance, **Fig 3**). Similar to the morning, movement was unaffected by recreational disturbance during the evening active bout at the population level, but between-individual variation increased along the recreational disturbance gradient (from conditional repeatability = 0.1 to conditional repeatability = 0.37, **Fig 3**). During the night resting bout, red deer showed little between-individual variation in their propensity to move (mean conditional repeatability = 0.1), which was also largely unaffected by the number of recreationists on nearby trails during the day time.

**Fig 3.**
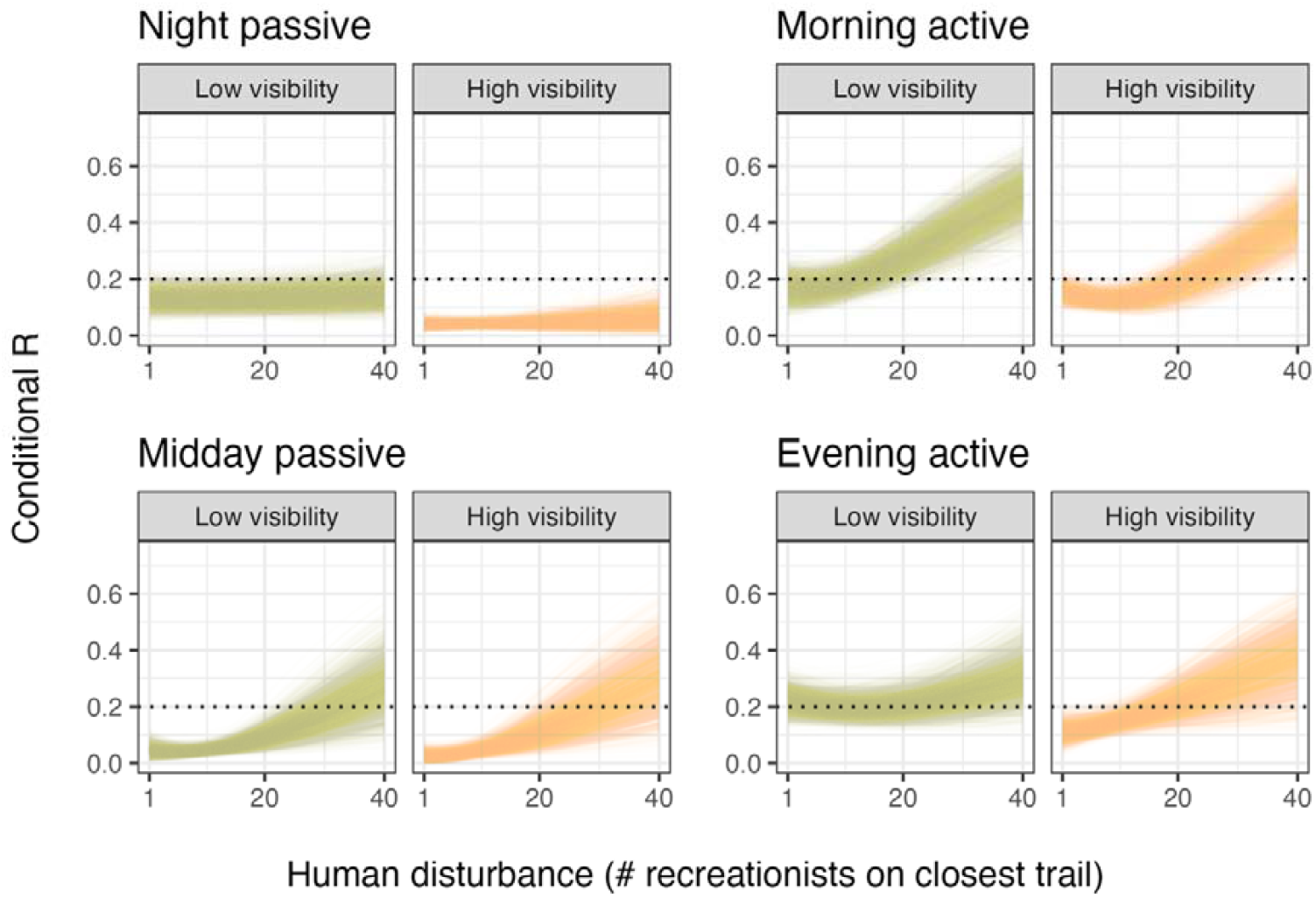
Conditional repeatability (i.e., variance standardized between individual variation) along a gradient of recreational disturbance under low and high visibility (10th and 90th percentile of visibility distribution). Lower conditional repeatability, suggests a similar average behavior of red deer or high within individual variability, while higher conditional repeatability suggests between-individual variation in behavioral expression. We added a dotted line at conditional repeatability = 0.2, which can be interpreted as a threshold beyond which repeatability can be interpreted as moderate to high.

## 4. Discussion

Red deer generally followed a crepuscular activity pattern with reduced movement activity in the middle of the day, when recreationists are active, and in the middle of the night, and instead primarily moved during the early morning and evening hours. There was little flexibility in diel activity patterns and all individuals followed the general population trend. Human disturbance such as recreational activity has previously been shown to impact red deer diel activity patterns, both in our study area but also in other regions, with more disturbed populations decreasing diurnal activity in comparison to less disturbed populations (Ensing et al., 2014; Kamler et al., 2007).

At the population level, movement activity of red deer was influenced by recreationists’ activity only during the middle of the day. Red deer responded to human disturbance by adjusting hourly step length, supporting our first hypothesis. We however did not find support for our first prediction, that red deer increase movement in response to disturbance. Rather, red deer responded with decreased movements (i.e., freeze) in the middle of the day, especially when in open vegetation. For example, while on the population level, movement during the middle of the day was generally low in dense vegetation (∼15 m/hr), movements in open vegetation decreased from ∼40 m/hr to ∼ 10 m/hr when deer were in proximity of a trail with many recreationists. A possible reason for why in the middle of the day deer do not increase movement rates to escape from recreationists is the very dense trail network providing no immediate refuge from human disturbance in case of a flight response, especially since recreational activities are also highest in the middle of the day (Neumann et al., 2010, Olson et al., 2018, Ordiz et al., 2019). Instead, red deer may opt to stay put on days with many recreational activities and instead ruminate. African dryland ungulates have been shown to increase ruminating time in the middle of the day as a response to decreased foraging activity, incurred by extreme heat (Berry et al., 2024). For red deer, we also found that deer shifted movements away from the hottest hours of the day and instead moved more during the morning, evening, and night, on days with high maximum daily temperature. At the population level, red deer did not respond to recreationists in the morning, evening or during the night. This could be because even on days with many recreationists in the park, disturbance is concentrated during the middle of the day. These results would be in line with the risk allocation hypothesis (Ferrari et al., 2009) and our second prediction and shows that red deer have highly nuanced responses to recreational activity, depending on the time of day and behavioral state. Whether total feeding time is reduced on days with high human disturbance or whether red deer shift activity to other times of day should be evaluated in future studies. However, the absence of an effect of recreation at the population level masked underlying individual differences in responsiveness to disturbance, especially during the morning and evening activity bouts.

In line with our second hypothesis, increasing human disturbance, measured as the number of recreationists on the trail closest to a deer, promoted between-individual variation in movement. This observation was contrary to the experimental literature, suggesting that animals might conform to a common phenotype under high risk while expressing greater variability under undisturbed conditions (Dingemanse et al., 2009). Individual variation in plasticity towards recreational disturbance was expressed during the morning, midday, and evening, on days and in areas with > 20 visitors on the closest trail and when deer moved through areas with high visibility and little concealing vegetation cover (supporting our prediction). During the morning activity bout, red deer forage preferably in such open areas and our results suggest that individual red deer solved the trade-off between foraging and perceived risk differently. For example, along a gradient of 1 to 40 recreationists on the closest trail, some individuals reacted strongly by increasing their hourly movement (e.g., from 300m/hr to 740m/hr) while others did not adjust movements. A contrasting behavioral response to recreationists was observed during the middle of the day. Again, between-individual variation increased with the number of recreationists on nearby trails but rather than increasing, red deer decreased movements in response to recreationists to varying degrees. Rapid or long-distance movements likely decrease the immediate time spent actively foraging, therefore, individuals that respond strongly likely incur a foraging cost. These ideas are in line with the pace-of-life syndrome theory, which predicts that individual variation in responsiveness towards risk incurs a risk-benefit trade-off where less fearful individuals are better at acquiring resources which leads to earlier reproduction and higher fecundity as compared to more fearful individuals (Réale et al., 2010). Individual variation in movement resulting from individuality in risk perception may thus be expected to translate into individual variation in resource acquisition, body mass, reproductive output per attempt, and survival (Biro and Stamps, 2008). However, evidence for the pace-of-life syndrome is scarce, in particular in natural systems (Royauté et al., 2018). It is unclear whether responsiveness translated into fitness costs in our population because we did not monitor reproductive success.

### Evolutionary implications

Following evolutionary theory, when behavioral trait variation is determined by genetic variation (i.e., heritability), natural or artificial selection for certain behavioral types, where individuals of a given behavioral expression survive better or reproduce more, can lead to shifts in behavioral trait composition over only a few generations (Lynch & Walsh 1998). There is good evidence that behavior is generally heritable (Dochterman et al. 2019). Though to-date rarely investigated, initial studies suggest a moderate heritability of routine movement behaviors such as average movement rate or daily movement speed in ungulates (mule deer and roe deer, respectively, Bonar et al., 2025; Gervais et al., 2020), opening the possibility for natural or artificial selection on these traits (Darimont et al., 2009; Leclerc et al., 2017). For example, hunting can lead to human-induced selection on behavioral traits and on individual plasticity to perceived risk (Ciuti et al., 2012; Madden and Whiteside, 2014; Otto, 2018). In fact, Ciuti et al., (2012) showed that elk differed in both, the average movement rate and in movement rate plasticity to human disturbance, and that hunters disproportionately harvested individuals with a higher movement rate especially when the risk of encountering humans was high. Therefore, elk with low behavioral plasticity incurred a survival cost (Ciuti et al., 2012; Thurfjell et al., 2017). Our results corroborate individual variation in the responsiveness to humans in red deer, in a system where hunting has been largely absent for the past 25 years. Responsiveness to recreational human disturbance, i.e., longer movements during morning hours when deer forage, therefore does not come with a survival benefit but incurs a foraging cost. It is therefore conceivable that natural selection, in the absence of or under limited hunter selection, increases the proportion of less reactive deer in the population. However, to support this idea future studies are needed to a) compare individual responsiveness to humans in hunted and non-hunted populations, b) establish whether fearfulness of humans is a heritable trait in wildlife, and c) whether there is generalizable evidence that hunters select for individuals with more conspicuous movement behavior.

### Methodological advancement

Pseudoreplication is a common pitfall when studying between-individual variation and is of particular concern in inherently autocorrelated movement data (Niemelä and Dingemanse, 2017). Pseudoreplication occurs when repeated measures of an individual are falsely treated as independent, e.g., when sequential hourly intervals are included as independent measurements in a model although the behavioral state of the animal precluded a change in behavior, here hourly movement distance (e.g., during resting or foraging on an aggregated food resource over sequential time steps). Including such non-independent data points can inflate between-individual variation because the individual supposedly shows a consistent response to the same environment, while in reality its behavioral state did not change. For movement data, repeated hourly steps of inactivity or course spatial resolution of environmental covariates, not matching the resolution of movement can lead to pseudoreplication (Hertel et al., 2020). By concentrating on specific hours of the day we could ascertain that repeated individual measures every 24hrs were truly independent. We advise that future studies of between-individual differences in animal movements pay attention to sources of pseudoreplication in their data, especially when stationary periods, such as resting, cannot be filtered out effectively, as was the case in our dataset.

## 5. Conclusion

We found great individual variation in behavioral plasticity towards recreational disturbance in a largely non-hunted red deer population. In particular, between-individual variation in movement behavior emerged at higher levels of perceived risk while individuals conformed to a common behavioral phenotype under low perceived risk from recreational activity. Some individuals responded with long movement distances to many recreationists in their vicinity, whereas others responded little. Whether responsive individuals pay a foraging cost or unresponsive individuals are more likely to be harvested in the BFNP hunting zone as well as outside the park should be subject of future research.

