## Supplementary material for "Red deer individual landscapes of fear in response to human recreation"

*Running header: Individual fear landscapes*

**Individual fear landscapes of red deer towards human recreation in a national park.**

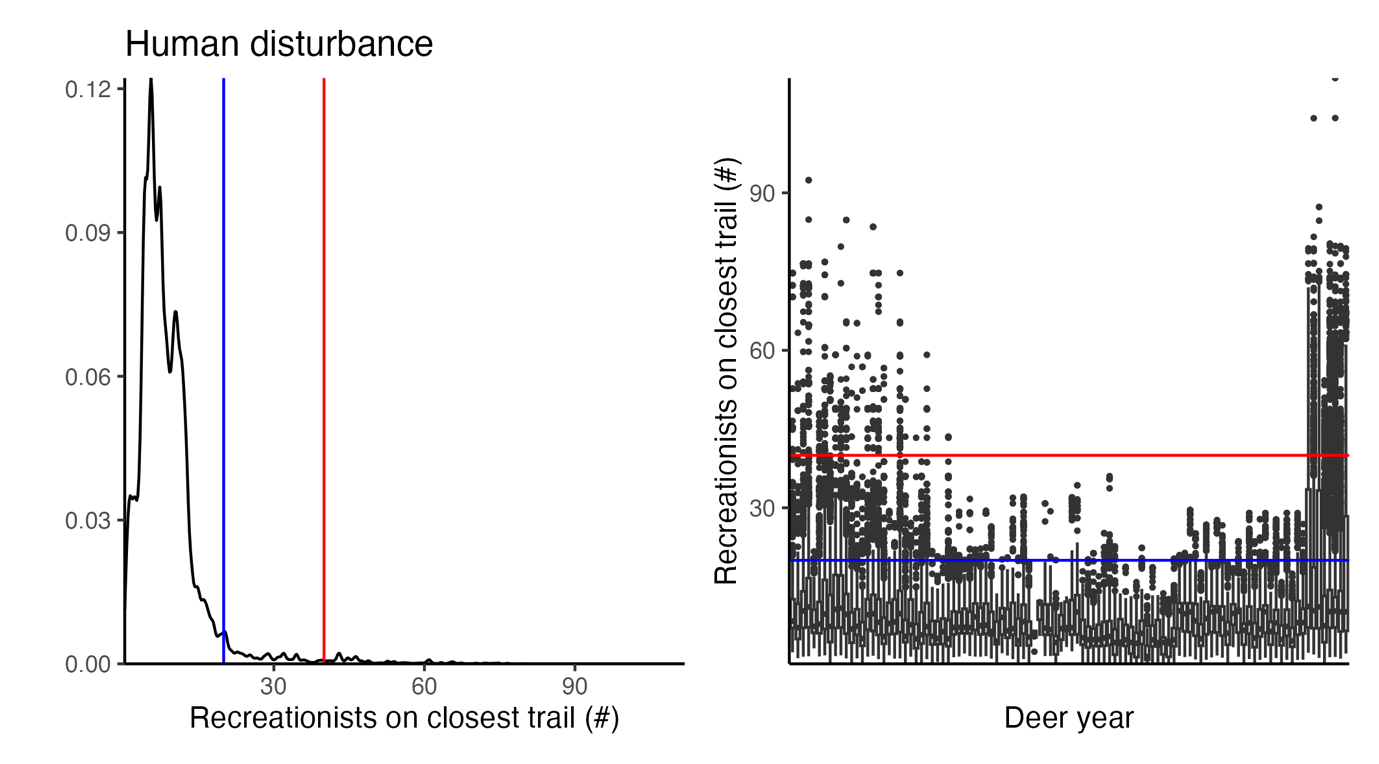

**Figure S1.** Distribution of human disturbance, measured as the predicted number of recreationists on the trail closest to a red deer GPS location. Most red deer locations were recorded on days and in areas of low human disturbance (3^rd^ quartile = 11 visitors), however almost all red deer experienced days with around 20 visitors in their vicinity (blue line). About half of the population experienced 40 visitors or more in their vicinity (red line).

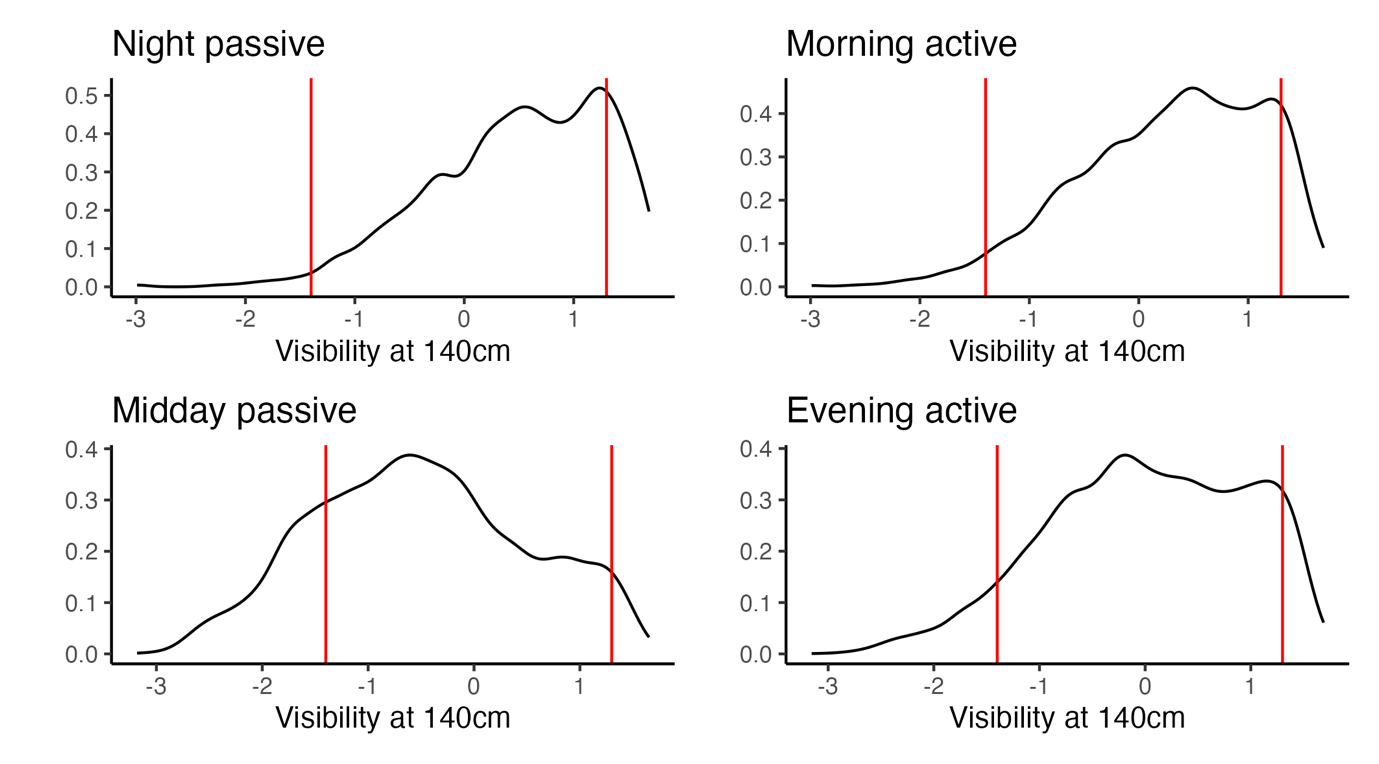

**Figure S2.** Distribution of visibility at 140cm height measured at GPS locations of red deer during the night passive, morning active midday passive, or evening active bout. Red lines indicate the 15^th^ and 85^th^ percentile of the overall occupied vegetation distribution. Red deer is found in denser vegetation during the middle of the day, when humans are most active in the park.

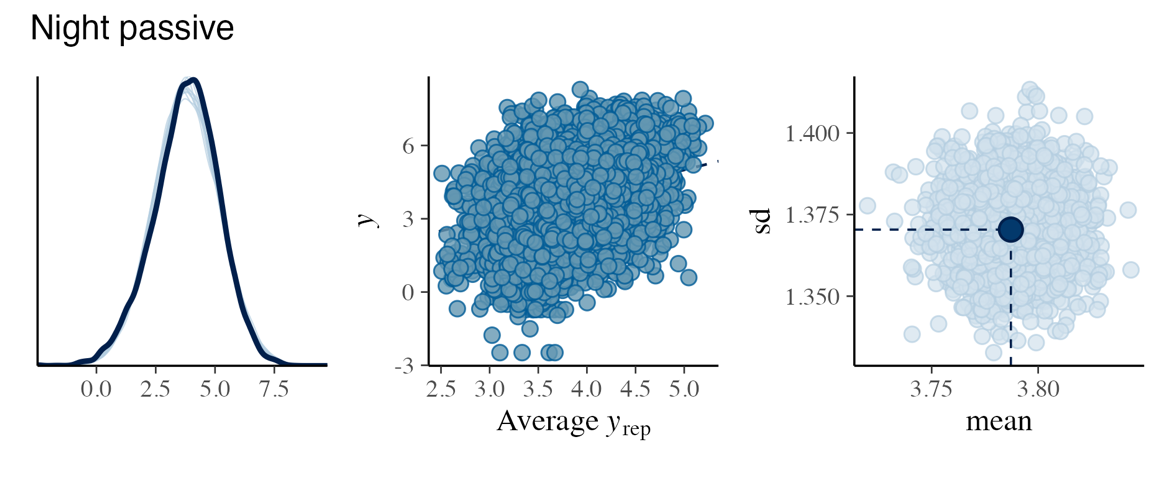

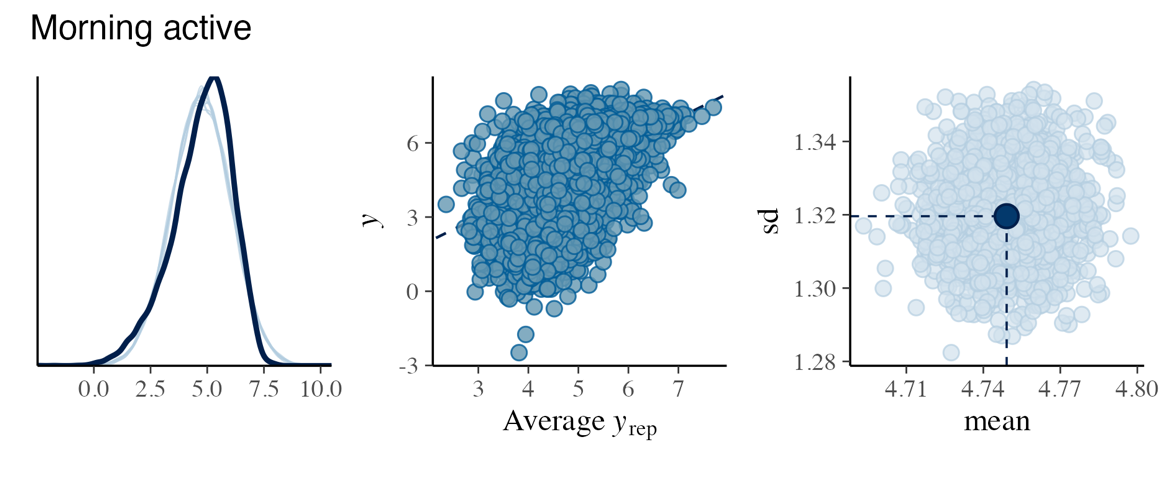

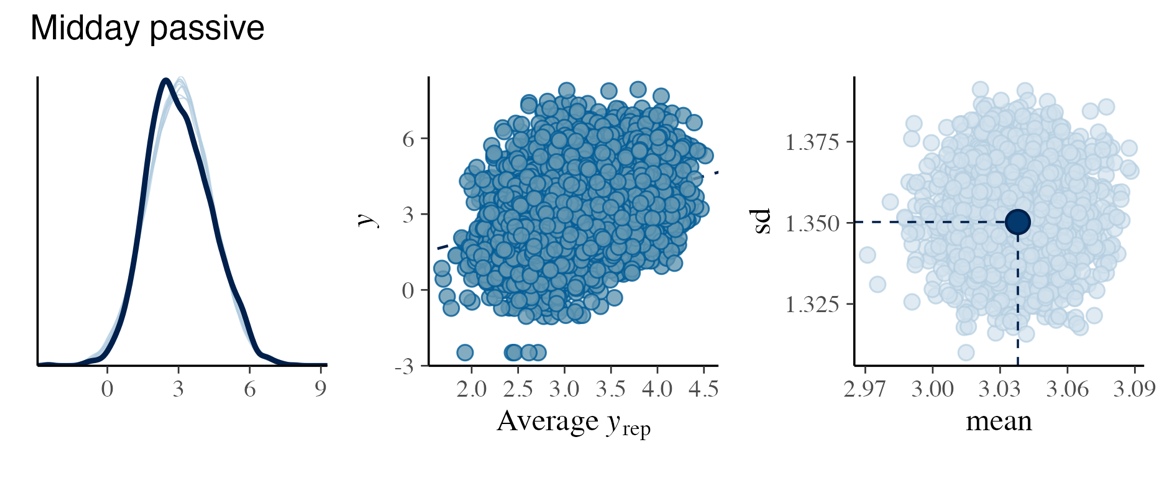

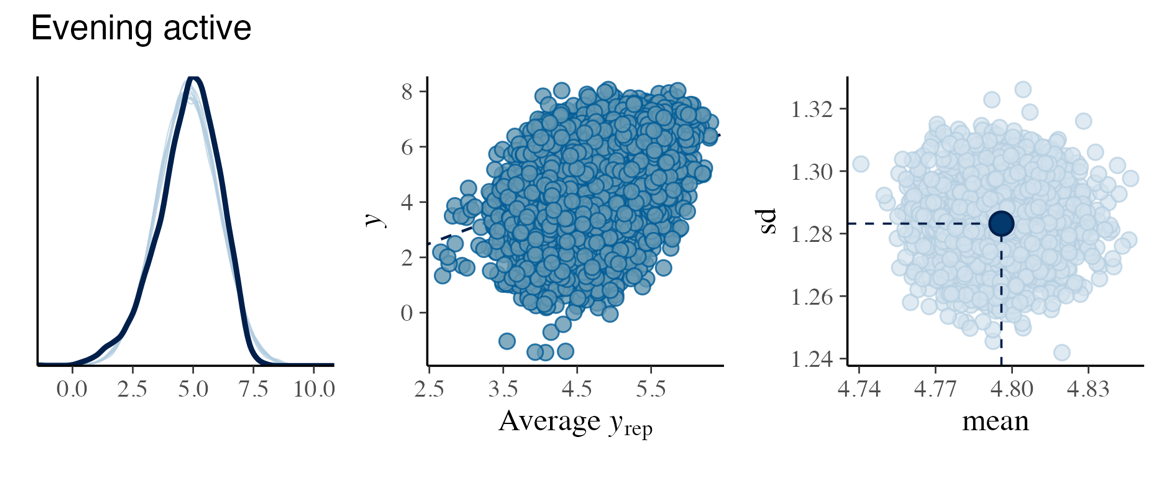

**Figure S3.** Posterior predictive checks for the four random regression models. Plots comparing the observed response variable 𝑦 (thick line) to simulated datasets 𝑦𝑟𝑒𝑝. (thin lines) from the posterior predictive distribution. The closer the two match, the better the model performs in reproducing the underlying distribution.
